# Differential splicing of ANP32A in birds alters its ability to stimulate RNA synthesis by restricted influenza polymerase

**DOI:** 10.1101/274613

**Authors:** Steven F. Baker, Mitchell P. Ledwith, Andrew Mehle

## Abstract

Adaptation of viruses to their host can result in specialization and a restricted host range. Species-specific polymorphisms in the influenza virus polymerase restrict its host range during transmission from birds to mammals. ANP32A was recently been identified as a cellular co-factor impacting polymerase adaption and activity. Avian influenza polymerases require ANP32A containing an insertion resulting from an exon duplication uniquely encoded in birds. Here we find that natural splice variants surrounding this exon create avian ANP32A proteins with distinct effects on polymerase activity. We demonstrate species-independent direct interactions between all ANP32A variants and the PB2 polymerase subunit. This interaction is enhanced in the presence of viral genomic RNA. In contrast, only avian ANP32A restored ribonucleoprotein complex assembly for a restricted polymerase by enhancing RNA synthesis. Our data suggest that ANP32A splicing variation amongst birds differentially impacts viral replication, polymerase adaption, and the potential of avian hosts to be reservoirs.

## Introduction

Influenza A viruses circulate in diverse host species. Wild aquatic waterfowl are the natural viral reservoir, and zoonoses can either occur directly from birds or through an intermediate host such as swine. Ecological overlap between the major hosts – birds, swine, and humans – creates repeated opportunities for cross-species virus transmission, yet only a minor fraction of these are successful. Influenza virus must overcome multiple biological barriers for successful cross-species transmission. The viral polymerase is a major determinant of host range (Almond, 1977; Subbarao et al., 1993). Avian-origin polymerases function efficiently in avian cells, but their activity is heavily restricted in human cells (Labadie et al., 2007; Mehle and Doudna, 2008). Restricted polymerases rapidly evolve adaptive mutations enabling efficient function as viruses jump from avian to mammalian hosts.

The influenza polymerase is a heterotrimeric enzyme composed of the subunits PB2, PB1, and PA. The polymerase assembles with viral RNA encapsidated by oligomeric nucleoprotein (NP) to form the viral ribonucleoprotein (vRNP) complex, the minimal unit required for viral transcription and replication (reviewed in (te Velthuis and Fodor, 2016)). The polymerase transcribes viral mRNAs via cap-snatching and replicates the minus-sense genomic vRNA through a plus-sense cRNA intermediate. Genome replication requires a *trans*-acting or - activating polymerase. Avian-origin polymerases are restricted in mammalian hosts with defects in both replication and transcription (Mehle and Doudna, 2008). The PB2 subunit has long been recognized as a main determinant of species-specific polymerase activity and host range (Almond, 1977; Labadie et al., 2007; Mehle and Doudna, 2008; Subbarao et al., 1993). The prototypical adaptive mutation in the PB2 subunit occurs at amino acid 627 located within the eponymous 627 domain, where an avian-signature glutamic acid is changed to a mammalian-signature lysine (Subbarao et al., 1993). Adaptive mutations in PB2 are associated with increased replication, pathogenicity, and transmission of avian-origin viruses in mammalian hosts. Recent structural analyses have revealed that portions of PB2, including the 627 domain, remain solvent exposed in the holoenzyme and undergo large-scale conformational reorganization depending on whether the polymerase is engaged in replication or transcription (te Velthuis and Fodor, 2016). These data raise the possibility that adaptive mutations in PB2 may be important for intra- or inter-molecular protein:protein interactions, which may also facilitate the conformational changes of the 627 domain.

Viral polymerase activity during infection is regulated by both essential host co-factors as well as restriction factors that antagonize function (Kirui et al., 2016a). Acidic nuclear phosphoprotein 32 family member A (ANP32A, pp32) associates with the influenza A virus polymerase and stimulates vRNA synthesis from a cRNA template *in vitro* (Bradel-Tretheway et al., 2011; Sugiyama et al., 2015). ANP32A has been implicated in diverse cellular roles including cell proliferation and cancer, apoptosis, transcriptional control, mRNA trafficking, and inhibition of protein phosphatase 2A (Reilly et al., 2014). More recently, ANP32A has been shown to impact the host range of influenza virus as a species-specific co-factor of the viral polymerase (Long et al., 2016). The restriction of avian-origin polymerases in mammalian cells is overcome by expressing avian ANP32A in these cells. Compared to mammalian ANP32A, which does not enhance avian polymerase activity, ANP32A encoded by most *Aves* species have a partial duplication of exon four resulting in an insertion between the N- and C-terminal domains. This insertion is necessary and sufficient to enable ANP32A to rescue restricted avian polymerases in mammalian cells (Long et al., 2016). While the genetic evidence strongly implicates ANP32A as a host-range factor, it remains unclear how ANP32A stimulates polymerase activity and how avian ANP32A selectively rescues restricted avian polymerases.

Here we dissect the processes by which ANP32A engages the viral polymerase and impacts its function. We identify naturally occurring splice variants of ANP32A in avian species that reduce the size of the repeat insertion from 33 to 29 amino acids, removing a SUMO interaction motif (SIM)-like sequence located upstream of the repeat. Both full-length chicken ANP32A (chANP32A_33_) and the splice isoform lacking the SIM (chANP32A_29_) rescue activity of a restricted polymerase, with chANP32A_33_ exhibiting a more potent phenotype. Expression of chANP32A_29_ in human cells is sufficient to increase replication of an avian-style virus. We show that ANP32A interacts with the viral polymerase, binding directly to the 627 domain. The presence of viral genomic RNA enhanced interactions between the polymerase and ANP32A. However, binding between ANP32A and the polymerase was not species-specific and was unaffected by the identity of PB2 residue 627. By contrast, our data reveal that chANP32A_29_ functions in a species-specific fashion to stimulate the intrinsic enzymatic activity of a restricted avian polymerase and alleviate defects in RNP assembly. Together, these data elucidate how chANP32A rescues polymerase activity and show that ANP32A splice variants present in birds affect its potency, with potential impacts on influenza adaptation and replication in these different hosts.

## Results

### Differential splicing creates three predominant isoforms of avian ANP32A with differing impacts on avian influenza polymerase activity

Species-specific differences in ANP32A impact its ability to function as a co-factor for the influenza virus polymerase (Domingues and Hale, 2017; Long et al., 2016; Sugiyama et al., 2015). We therefore analyzed ANP32A expression in diverse avian species. ANP32A contains a conserved N-terminus with leucine-rich repeats and a largely disordered low-complexity acidic region at the C-terminus (Reilly et al., 2014). Most avian *ANP32A* encode a partial duplication and insertion of exon 4 that repeats a portion of the leucine-rich repeat capping motif in the expressed protein (**Fig. 1A**). This duplication is absent in mammals and the avian Palaeognathae clade containing ostriches and tinamous. Analysis of RNA-seq datasets revealed alternative splicing surrounding the duplicated exons (**Fig. 1A-B**). The chANP32A that was originally shown to enhance polymerase activity contained a 33 amino acid insertion. Inspection of intron-spanning reads revealed alternative splicing to a downstream splice acceptor site to create chANP32A_29_. chANP32A_29_ lacks four hydrophobic residues from the N-terminus of the repeat that compose the SIM present in the 33 amino acid insert (Domingues and Hale, 2017). The ratio of transcripts encoding 33 or 29 amino acid inserts varied across species (**Fig. 1C; Table S1**): pigeons (*Columba livia*) express almost exclusively _ANP32A33_, chickens (*Gallus gallus*) express ∼3 fold more ANP32A_33_ transcripts than _ANP32A29_, and starlings (*Sturnus vulgaris*) express almost exclusively ANP32A_29_ transcripts. Intriguingly, examples were found in transcripts from many avian species where the duplicated exon was skipped altogether to create a human-style ANP32A. This was most pronounced in the swan goose (*Anser cygnoides*) where >70% of all transcripts skipped the duplicated exon. ANP32A splicing patterns were unaffected by influenza virus infection in chickens and did not differ between inbred lines that are susceptible or partially resistant to influenza A virus (**Table S1**). Further, there was no obvious difference in ANP32A splicing in chicken cells when IFN signaling was activated or inhibited, or when they were infected with influenza virus (**Table S1**).

**Figure 1.**
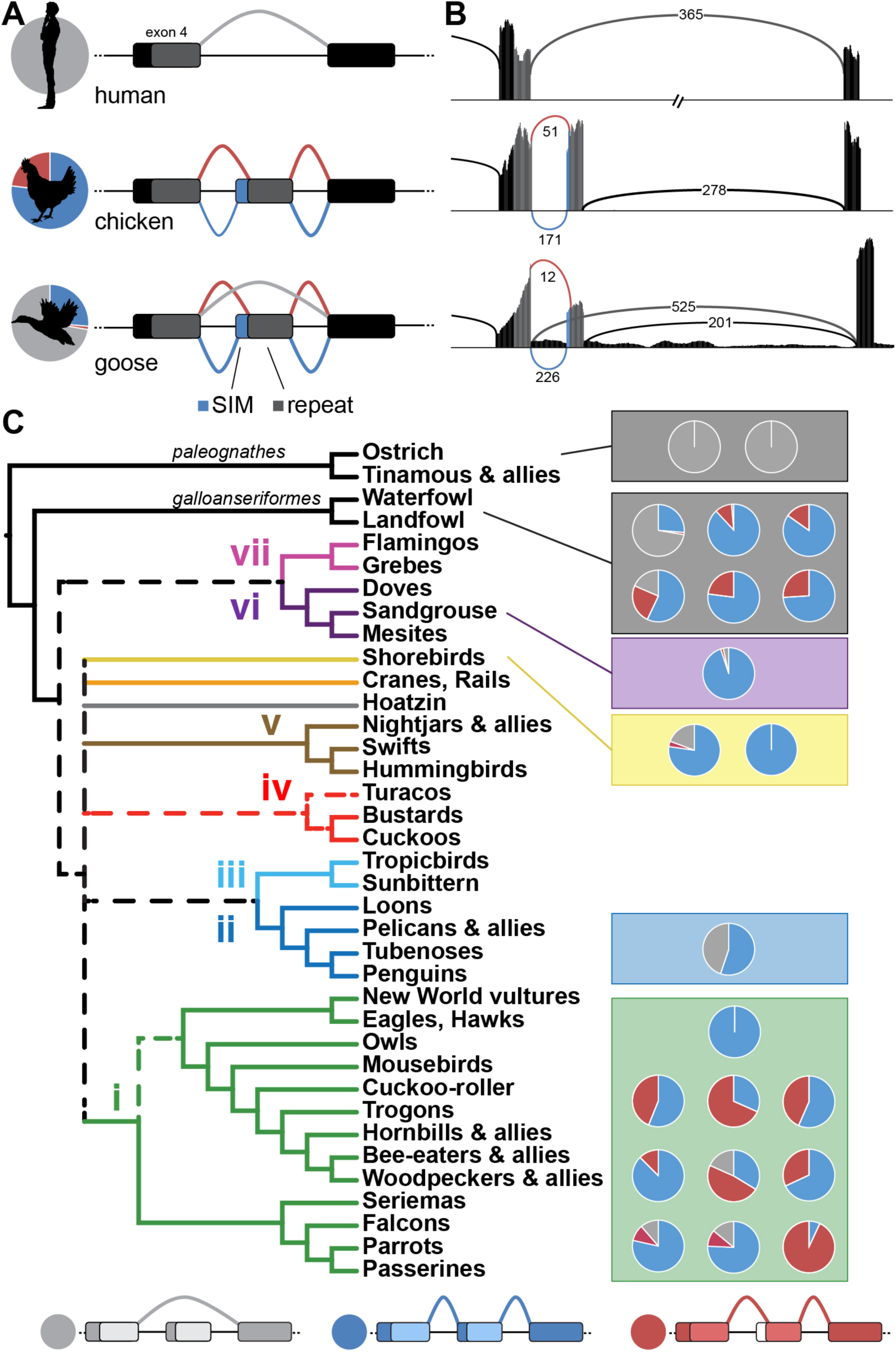
Natural variation in ANP32A splicing patterns in *Aves*. Duplication and insertion of *ANP32A* exon 4 results in three major splice isoforms in birds. (**A**) All human ANP32A transcripts lack the exon duplication (light gray). Schematics of chicken and goose transcripts show splicing upstream to capture coding sequence for the SIM (blue), splicing downstream to omit the SIM (red), or in some cases skipping the repeated exon altogether to create a mammalian-like transcript (light gray). The relative abundance of each splice isoform in RNA-seq data is indicated by the pie charts. SIM; SUMO-interacting motif. (**B**) Sashimi plots of *ANP32A* corresponding to examples in (A) and colored similarly. The abundance of each splice variant is indicated on the lines corresponding to the intron-spanning reads. (**C**) ANP32A splice patterns in diverse bird species. *Aves* consensus phylogeny where paleognathes are the major outgroup and the seven major groups are indicated (based on (Reddy et al., 2017)). Pie charts represent relative transcript abundance from RNA-seq datasets (listed in **Table S1**).

To assess the impact of ANP32A splice variants, we performed polymerase activity assays in human or avian cells by reconstituting RNP complexes with a vRNA-like firefly luciferase reporter gene. Avian-style polymerases (PB2 E627) were heavily restricted in human cells, whereas co-expression of chANP32A_33_ restored activity in a dose-dependent fashion (**Fig. 2A**), consistent with earlier reports (Long et al., 2016). The enhancing activity of chANP32A_33_ was extremely potent, increasing polymerase activity even when its expression was below the limit of detection in our western blots. Co-expressing chANP32A_33_ had minimal effects on a human polymerase (PB2 K627) in human cells. Blotting confirmed that the increased polymerase activity was independent of changes in viral protein levels.

**Figure 2.**
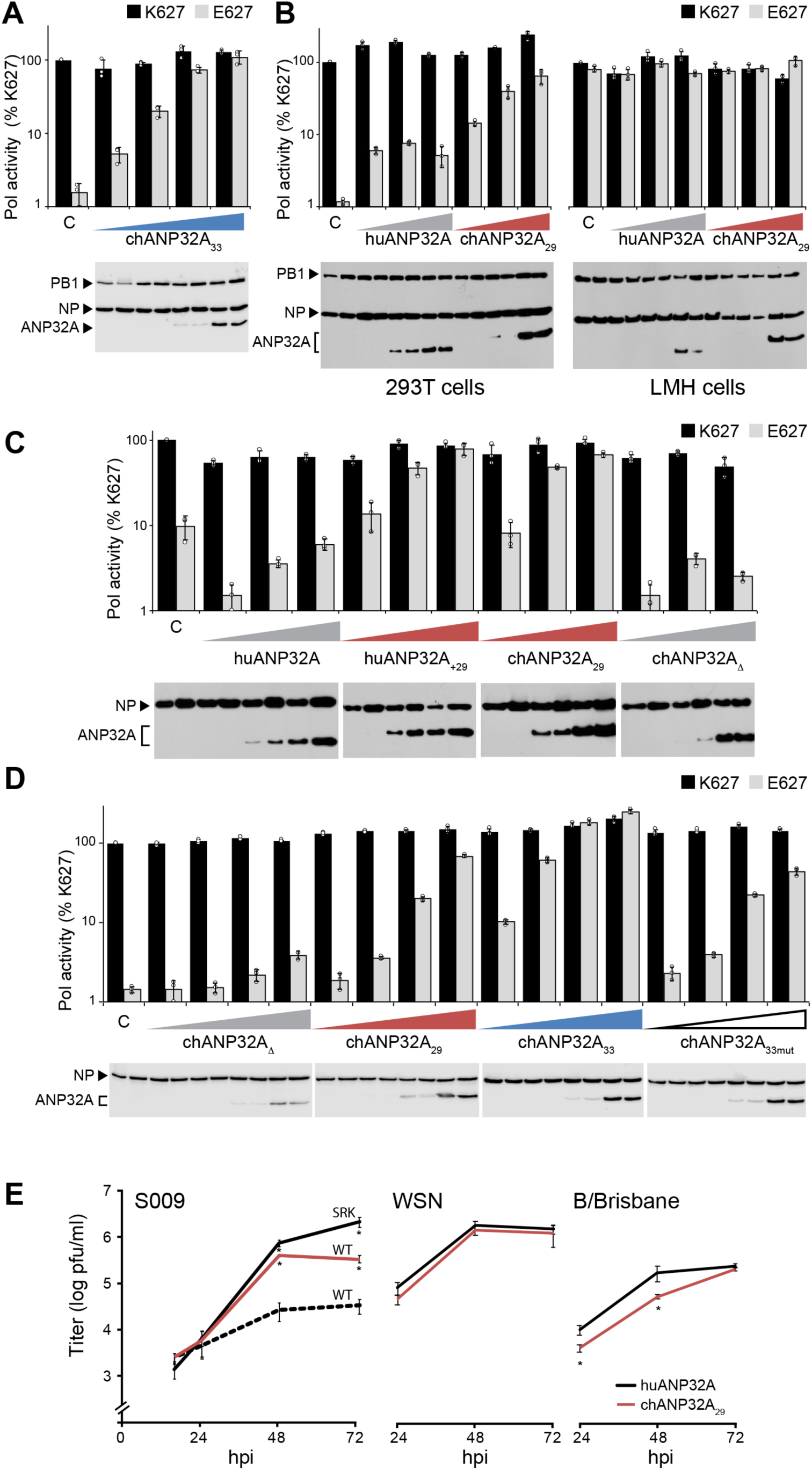
Avian ANP32A29 is sufficient to restore species-restricted avian polymerase activity and replication. Polymerase activity of both human influenza polymerase (PB2 K627) and avian-style polymerase (PB2 E627) was measured in the presence of increasing concentrations of the indicated ANP32A proteins. Protein expression was assessed via western blot. (**A**) ANP32A33 selectively rescues restricted avian polymerase activity in human 293T cells. (**B**) ANP32A29 is sufficient to restore polymerase activity in human cells (*left*), but does not significantly impact human viral polymerase in human cells or either polymerase in avian LMH cells (*right*). (**C**) Insertion of the avian 29 amino repeat into huANP32A (huANP32A+29) enhances activity whereas deletion of the repeat in chANP32A (chANP32AΔ) disables its function. (**D**) chANP32A33 is the most potent enhancer of avian polymerase activity in human cells compared to chANP32A29 or chANP32A33mut. For all assays, polymerase activity was normalized to an internal control and compared to PB2 K627 polymerase in the absence of ANP32A. Data are shown as average of *n* = 3 +/- standard deviation derived from representative results of at least 3 independent experiments. (**E**) Replication kinetics of influenza virus encoding the avian S009 RNP (WT) or a human human-adapted mutant (SRK; G590S, Q591R, E672K), WSN, or B/Brisbane. A549 cells stably expressing ANP32A were infected (MOI = 0.1) and viral titers were determined at the indicated time points. Data are shown as average of *n* = 3 +/- standard deviation. *p<0.05 one-way ANOVA when compared to WT A549 cells.

Given the prevalence of ANP32A_29_ in relevant avian hosts, we assessed the capacity of this splice variant to support avian influenza virus polymerase activity. Expressing chANP32A_29_ in restrictive human cells rescued the activity of an avian-style polymerase in a dose-dependent manner (**Fig. 2B**). chANP32A_29_ did not significantly impact the activity of a human influenza virus polymerase in human cells, or either polymerases in avian cells. Expressing human ANP32A (huANP32A) exhibited only minimal effects on both polymerases. We created ANP32A domain swap hybrids to determine if this shortened insert is sufficient to enhance polymerase activity (**Fig. 2C**). Inserting the 29 amino acid repeat into huANP32A (huANP32A_+29_) conferred enhancing activity to this hybrid protein, which supported avian polymerase activity at levels comparable to chANP32A_29_. Conversely, removing the 29 amino acid insert from chANP32A (chANP32AΔ) eliminated its enhancing activity. The four amino acids that are absent in the insert of ANP32A_29_ have been identified as a SIM-like sequence important for the pro-viral activity of chANP32A_33_ (Domingues and Hale, 2017). We thus tested the relative activity of chANP32A variants (**Fig. 2D and Supplemental Figure 1**). Both splice variants of chANP32A increased activity of an avian polymerase in human cells, although the dose-dependent assay revealed that chANP32A_33_ is over 10 fold more effective than a similar amount of chANP32A_29_. This difference in activity was dependent upon the SIM sequence, as simply restoring the insert length to 33 amino acids by adding four glycine residues on the N-terminus to create ANP32A_33mut_ did not increase activity above that of chANP32A_29_ (**Fig. 2D**).

To determine if the observed changes in polymerase activity increase viral replication, we performed multi-cycle replication assays in human cells stably expressing huANP32A or chANP32A_29_. The avian influenza virus isolate S009 possesses glutamic acid at residue 627 in PB2 and is restricted in human cells (Mehle and Doudna, 2009). Virus growth kinetics showed that expression of chANP32A_29_ increased replication of the restricted S009 WT approximately 10-fold (**Fig. 2E**). Cells were also infected with an S009 mutant encoding a fully humanized PB2 with a serine at position 590, an arginine at position 591, and a lysine at position 627 (S009 SRK) (Mehle and Doudna, 2009). As expected, the mutations present in S009 SRK overcame the restriction phenotype in human cells. Restoring replication by expressing chANP32A_29_ in the target cells or by introducing adaptive mutations into PB2 produced similar viral titers at early time points, although the adapted virus reproducibly achieved higher final titers. Infections were repeated with mammalian influenza A (A/WSN) and B (B/Brisbane) viruses (**Fig. 2E**). In these experiments, expressing ANP32A had no significant effect on WSN replication and only minor effects on replication at early times for B/Brisbane. These data suggest that chANP32A_33_ containing the SIM sequence is the more potent enhancer of avian polymerase activity while also revealing that the shorter 29 amino acid insert encoded by certain splice isoforms of avian ANP32A is sufficient for supporting polymerase activity and replication of an avian-style virus.

### ANP32A interacts directly with the 627 domain of PB2

The mechanism(s) by which chANP32A enhances activity of avian-origin polymerases remain obscure. We performed a series of experiments to dissect potential interactions between ANP32A and the viral RNP. The remainder of our experiments focused on chANP32A_29_ given that it contained a minimal insert that supported avian polymerases while lacking the SIM motif, allowing us to segregate SIM-related activities from the effects of the common repeat sequence. We tested interactions between PB2 and ANP32A in human cells using human and avian versions of both proteins (**Fig. 3**). To focus solely on the primary determinants of polymerase adaption, we used WSN PB2 variants that differed only at residue 627 to encode either the human-signature lysine or the avian-signature glutamic acid. Similarly, we utilized chANP32A_29_ or a variant where the repeat has been deleted (chANP32AΔ), which mimics the human version while controlling for the other 18 amino differences between bird and human homologs. Coimmunoprecipitations detected the known interaction between NP and PB2 but failed to detect stable binary interactions between ANP32A and PB2, regardless of which combination of human and avian proteins were used (**Fig. 3A**). By contrast, ANP32A specifically co-precipitated with PB2 when the subunits PB1 and PA were also present (**Fig. 3B**), suggesting that ANP32A preferentially interacts with the trimeric polymerase and in agreement with prior work (Bradel-Tretheway et al., 2011; Sugiyama et al., 2015). We confirmed these results in the context of infected cells where PB2 specifically co-precipitated ANP32A (**Fig. 3C**).

**Figure 3.**
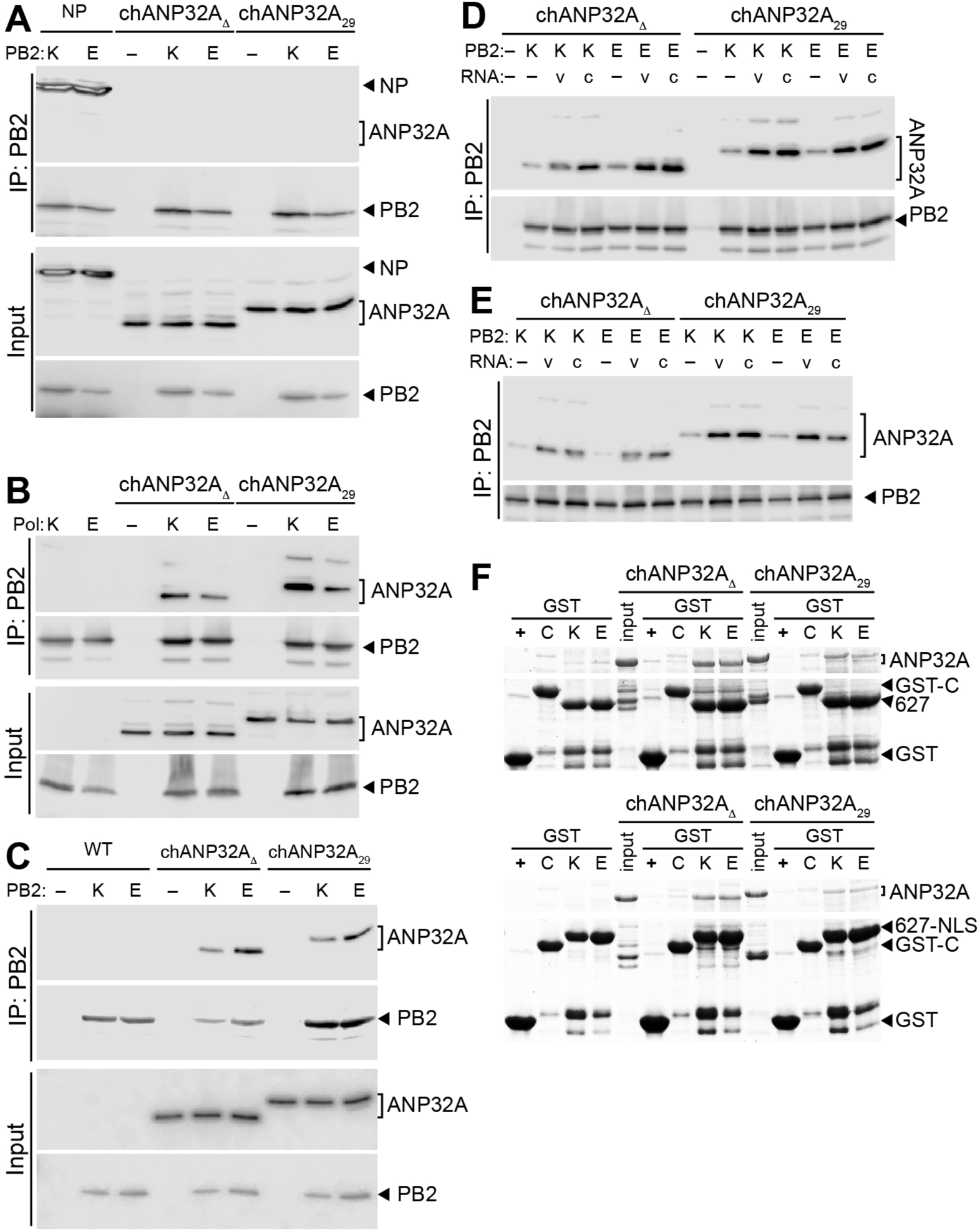
ANP32A binds directly to the PB2 627 domain in a species-independent fashion. (**A**) ANP32A does not stably interact with the PB2 subunit when it is expressed alone. PB2 K627 or E627 was immunoprecipitated (IP) from cells co-expressing ANP32A or the positive control NP. (**B**) Both chANP32A29 and the humanized chANP32AΔ interact with the polymerase trimer. Immunoprecipitations were performed from cells expressing ANP32A and the trimeric polymerase containing PB2 K627 or E627. (**C**) ANP32A interacts with the viral polymerase during infection. WT A549 cells and those stably expressing ANP32A were infected with PB2 K627 or PB2 E627 virus and PB2 was immunoprecipitated. (**D-E**) Genomic RNA increases PB2-ANP32A interactions. Immunoprecipitations were performed from cells expressing ANP32A, a vRNA (v) or cRNA (c) genomic segment, and the trimeric polymerase containing PB2 K627 or E627. Binding was measured with catalytically active polymerase (D) or an inactive PB1a mutant polymerase (E). For A-E all proteins were detected by western blot. (**F**) ANP32A binds directly to the PB2 627 domain. *In vitro* binding was measured between chANP32A variants and GST-tagged PB2 627 domain (top) or PB2 627-NLS (bottom) and visualized by Coomassie staining. GST alone (+) or an irrelevant GST fusion (C) were included as specificity controls.

Notably, there was no striking evidence for a species-specific interaction as both chANP32A_29_ and chANP32AΔ interacted equally with PB2 K627 and PB2 E627 polymerases. In addition, these results show that the SIM was not required for interactions between chANP32A and the polymerase. Thus, while chANP32A, but not huANP32A, rescues restricted avian polymerases and viruses in human cells (**Fig. 2**), the selectivity of this enhancement does not appear to occur at the level of protein:protein interactions.

The viral polymerase, and PB2 in particular, adopts multiple conformations depending on the function of the enzyme and co-factors present (te Velthuis and Fodor, 2016). We co-expressed vRNA or cRNA genomic templates with the polymerase and ANP32A to bias our protein interaction analyses and shift the polymerase towards an RNA-bound conformation, (**Fig. 3D**). NP was intentionally excluded to prevent genome replication, which would result in a mixture of vRNA and cRNA templates. As above, PB2 K627 and E627 co-precipitated both chANP32A_29_ and chANP32AΔ. Co-expressing either vRNA or cRNA significantly increased co-precipitation of chANP32A_29_ and chANP32AΔ. This increase was similar for plus-or minus-sense templates and human or avian PB2. To ensure that nascent products are not being produced, we repeated the interaction assays using a catalytically defective PB1 (PB1a). Genomic RNA increased interactions between the polymerase and ANP32A even when the polymerase was catalytically inactive (**Fig. 3E**). These findings suggest that ANP32A interacts more efficiently with RNA-bound conformations of the polymerase and these interactions do not require catalysis. The fact that this enhancement occurred with both cRNA and vRNA templates raises the possibility that ANP32A prefers a conformation poised for replication, as cRNA-bound polymerases do not normally perform transcription. Additionally, providing genomic RNA establishes distinct populations of *cis* and *trans* polymerases; the increase in ANP32A binding in the presence of genomic RNA may also reflect a preference for one of these populations.

Most adaptive mutations in avian polymerases arise within the conformationally dynamic PB2 627 and NLS domains (Manz et al., 2013; te Velthuis and Fodor, 2016). To evaluate if ANP32A interacts directly with the 627 domain of PB2, *in vitro* binding was tested using recombinant purified proteins. chANP32A_29_ and chANP32AΔ were captured by the PB2 627 domain, but not by an irrelevant control GST fusion protein or GST alone (**Fig. 3F, *upper***). Similar results were obtained using PB2 containing the 627 and NLS domains (**Fig. 3F, *lower***). Once more, these interactions were not dependent on the presence of the repeat insert in ANP32A, the SIM, or the PB2 variant used. We note that these binding assays were performed at steady state, and minor differences in binding kinetics may not be detectable via this approach. Together, our cell-based experiments and *in vitro* binding assays show that ANP32A directly binds the 627 domain of PB2, and that this interaction is enhanced when the polymerase adopts a template-bound conformation.

### The molecular defects of PB2 627E polymerase in human cells are nullified by chANP32A

Avian polymerases cannot efficiently assemble RNPs in mammalian cells (Mehle and Doudna, 2008). We thus sought to evaluate if RNP formation was affected by ANP32A (**Fig. 4A**). Polymerase, NP and vRNA were expressed in human cells and co-precipitation of PB2 by NP was used as a proxy for RNP formation. Avian-style polymerase exhibited the characteristic defect in RNP formation (**Fig. 4A**). Co-expressing chANP32A_29_ or chANP32AΔ in human cells did not alter RNP assembly for human polymerase complexes. By contrast, co-expressing chANP32A_29_ overcame assembly defects for avian-style polymerases as evidenced by a significant increase in PB2 co-immunoprecipitation. chANP32AΔ did not restore RNP formation, demonstrating specificity of the enhancement. These results demonstrate that the rescue phenotype observed in viral polymerase activity assays (**Fig. 2**) are in part due to the ability of chANP32A_29_ to enhance PB2 627E RNP formation.

**Figure 4.**
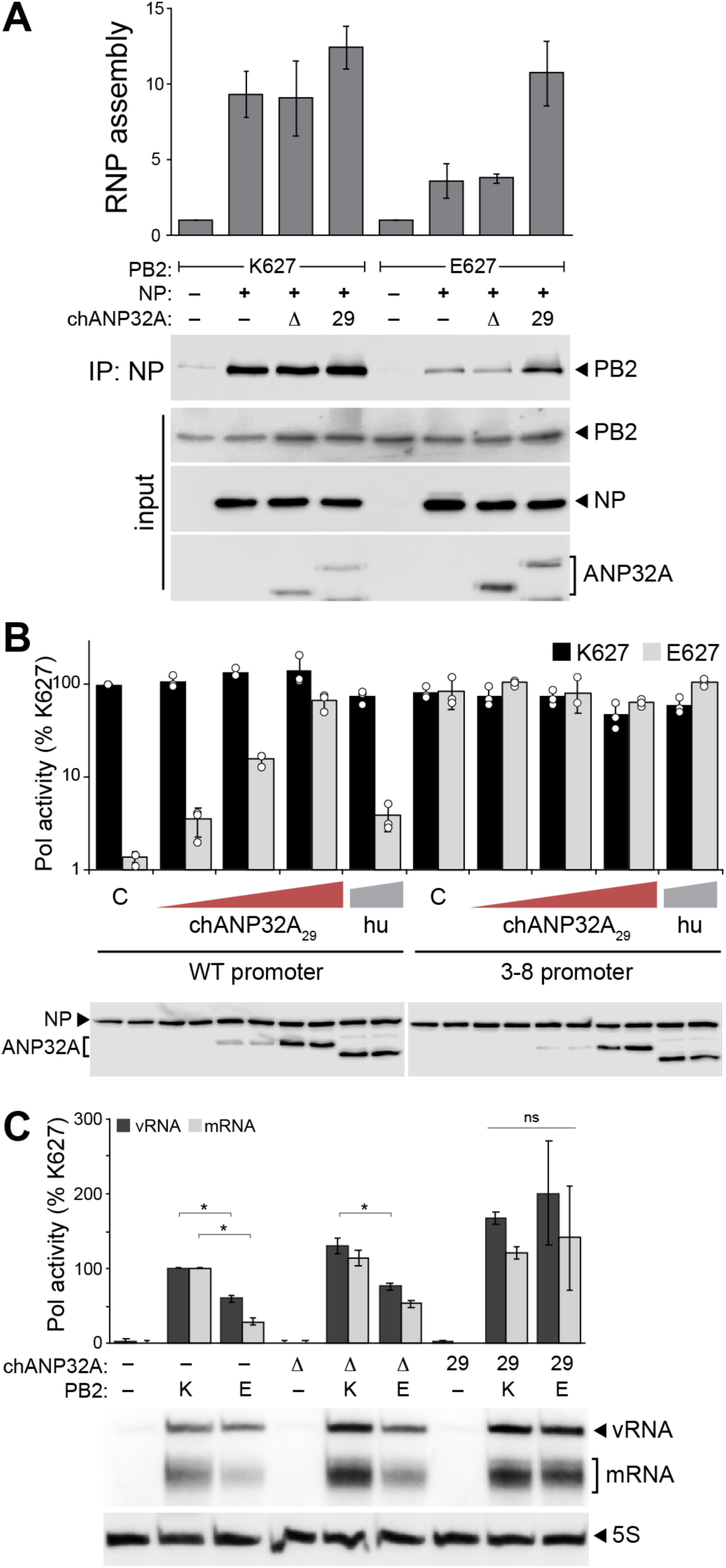
Molecular defects of a restricted avian polymerase are rescued by chANP32A_29_. (**A**) chANP32A_29_ enhances RNP formation for an avian-style polymerase. RNPs were reconstituted in human cells by co-expressing the indicated polymerase, NP, a genomic RNA and chANP32A variants. NP was immunoprecipitated and co-precipitating PB2 was measured as a proxy for RNP formation. Proteins were detected by western blot. RNP assembly was quantified from two independent assays, normalized to controls, and represented as the mean +/- standard deviation. **(B)** chANP32A_29_ functions redundantly with enhancing promoter mutations. Polymerase activity assays were conducted with the indicated ANP32A proteins and either WT or 3-8 mutant vRNA reporters. Proteins were detected by western blot. (**C**) chANP32A_29_ increases the enzymatic activity of a restricted polymerase independent of RNP formation. NP-independent polymerase activity on a micro gene vRNA template was measured by primer extension. chANP32A variants were co-expressed where indicated and PB2 was omitted as a negative control. mRNA and vRNA products were quantified by phosphorimaging from two independent experiments, normalized to PB2 K627 in the absence of ANP32A, and presented as mean +/-standard deviation. *p<0.05 one-way ANOVA.

Defects in RNP formation by restricted polymerases are proposed to result from decreased production or stabilization of replication products by the polymerase itself rather than changes to downstream events in the assembly process (Cauldwell et al., 2013; Paterson et al., 2014). Mutations in the 3’ vRNA promoter at positions 3 and 8 that increased RNA synthesis by the polymerase are also overcame restriction for avian polymerases (**Fig. 4B** and (Cauldwell et al., 2013; Paterson et al., 2014)). Co-expressing chANP32A_29_ with a 3-8 mutant vRNA reporter did not further increase polymerase activity. The absence of additive effects suggests that the promoter mutant and chANP32A_29_ alter polymerase activity at a similar step, possibly through the same pathway (**Fig. 4B**). We next eliminated NP and asked if ANP32A impacts the polymerase itself to change synthesis or accumulation of RNA products. Although the viral polymerase requires NP for synthesis of full-length genomic segments, NP is not necessary to replicate and transcribe short (∼76 nt or less) micro vRNA-like templates (Turrell et al., 2013). Human viral polymerase produced mRNA and vRNA products from a micro vRNA template, but an avian-style polymerase remained restricted with significant reductions in all RNAs (**Fig. 4C**). Co-expressing chANP32AΔ slightly increased synthesis of vRNA and mRNA, yet avian-style polymerase remained restricted. However, co-expressing chANP32A_29_ restored avian-style polymerase product accumulation to human polymerase levels. Together these data suggest that chANP32A_29_ is a minimal variant that overcomes defects in RNA synthesis by restricted avian-style polymerases, restoring RNA production resulting in RNP formation and subsequent viral gene expression.

## Discussion

Avian influenza virus polymerases are heavily restricted in mammalian cells. This is due in part to the species-specific dependence on the host factor ANP32A. An exon duplication and insertion present in most *Aves* species confers replication competence to influenza virus encoding an avian-style polymerase. Here we have shown that this duplicated exon is alternatively spliced in bird species creating ANP32A proteins with differential ability to support avian influenza polymerases. chANP32A_33_ containing a SIM sequence was the most potent isoform, and the most prevalent variant in common reservoir hosts such as ducks and chickens. The chANP32A_29_ variant, which lacks the SIM sequence, was sufficient to rescue avian polymerase function in mammalian cells. In contrast, transcripts that skipped the duplicated exon altogether produced chANP32AΔ that mimicked mammalian ANP32A and failed to rescue polymerase activity. ANP32A bound directly to the PB2 627 domain and binding in cells was enhanced by the presence of genomic RNA. PB2 and ANP32A interacted equivalently regardless of which ANP32A variant was tested or whether PB2 was avian-or human-origin. Species specificity of ANP32A only became apparent when investigating the intrinsic activity of the viral polymerase. chANP32A_29_ selectively enhanced production of RNA products by restricted avian-style polymerases. Together these data show that natural splice variants in ANP32A impact its ability to selectively enhance the production of RNA products by restricted polymerases.

The precise step(s) at which avian polymerases are restricted in mammals, and where ANP32A functions to alleviate restriction, is not well understood. Restricted polymerases express, localize, assemble into holoenzymes, and bind to genomic RNA similarly to functional polymerases (Mehle and Doudna, 2008; Nilsson et al., 2017). The use of promoter mutants to bypass restriction has also shown that restricted polymerases do not have defects in elongation (Crescenzo-Chaigne et al., 2002; Paterson et al., 2014). When restricted polymerases are purified from human cells they retain full activity in *in vitro* assays indicating that components in the cellular environment alter their activity (Paterson et al., 2014). Thus, restriction appears to occur at the earliest steps in the catalytic process, after template binding and before elongation. This is also the stage at which our data show enhanced ANP32A:polymerase interactions and species-specific enhancement of RNA synthesis (**Figs. 2, 4**). 3-8 promoter mutants can also function at this step to drive increased production of viral RNAs. Our demonstration that chANP32 showed no additive effects with the 3-8 promoter mutant provides further support that ANP32 impacts these early steps in RNA synthesis. This supports a model where ANP32A stimulates intrinsic activity of restricted polymerases by favoring a conformation poised for replication or by recruiting necessary *trans*-acting or -activating polymerases.

Our data raise the possibility that host-specific expression and splicing patterns of *ANP32A* apply distinct selective pressures on influenza virus. Ostrich *ANP32A* does not encode a duplicated exon four (Long et al., 2016). Instead, they express a human-like ANP32A, which likely explains why experimental infection of ostriches with an avian influenza virus resulted in acquisition of mammalian adaptive mutations in PB2 (Shinya et al., 2009; Yamada et al., 2010). Similar dynamics may have been in play during a highly pathogenic avian influenza outbreak at Qinghai Lake in 2005. Avian viruses isolated from this outbreak encoded the prototypical human adaptation PB2 E627K (Chen et al., 2005, 2006; Liu et al., 2005).

Bar-headed geese (*Anser indicus*) were the index species for this outbreak. RNA-seq analysis from the closely related swan goose (*Anser cygnoides domesticus*) revealed stark differences in ANP32A splicing patterns with >70% of transcripts skipping the repeated exon to express a human-like ANP32A (**Fig. 1, Table S1**). Our mechanistic data show that this pattern of ANP32A expression would pressure the emergence of human-signature adaptations in PB2. Thus, inherent host differences in ANP32A splicing transcripts have the potential to shape the evolution of PB2 in natural avian hosts and poise viruses for cross-species transmission by “pre-adapting” them for replication in mammals. In summary, we show that chANP32A isoforms differentially stimulate the enzymatic activity of restricted polymerases and posit that host-specific ANP32A splicing patterns affect replication within and transmission from avian reservoirs.

## Experimental Procedures

Detailed materials and methods descriptions can be found in Supplement Experimental Procedures.

### RNA sequencing analysis

HISAT2 was used to align RNA-seq data sets detailed in Table S1 to ANP32A genomic DNA assemblies from their respective species (Kim et al., 2015). Differential splicing events were visualized in IGV using sashimi plot function (Katz et al., 2015).

### Viruses and plasmids

Viruses and plasmid clones were derived from A/WSN/33 (H1N1; WSN) and A/green-winged teal/Ohio/175/1986 (H2N1; S009) (Mehle and Doudna, 2008, 2009). Recombinant viruses were produced using the influenza virus reverse genetics system (Neumann et al., 2005). Viral stocks were confirmed by sequencing and titered by plaque assay. Multi-cycle growth curves were performed in A549 cells and samples collected at the indicated time points were titered by plaque assay.

Mammalian and bacterial expression vectors were created for chANP32A_29_ (synthesized by IDT) or huANP32A (DNASU Plasmid Repository HsCD00042415). ANP32A variants were created by PCR-based mutagenesis. Influenza virus reporter gene plasmids and micro gene constructs were previously described (Kirui et al., 2016b).

### Polymerase activity assays, co-immunoprecipitations and in vitro binding

RNP complexes were reconstituted by expressing PB2, PB1, PA, NP, vNA-Luc reporters, ANP32A where indicated, and a constitutively expressed Renilla luciferase control (Mehle and Doudna, 2008). Firefly luciferase was normalized to the internal Renilla control. Protein levels in lysate were evaluated by Western blotting. Co-immunoprecipitations were performed in 293T cells under similar conditions of RNP reconstitution using FLAG-tagged PB2. PB2 was immunoprecipitated with anti-FLAG agarose resin (M2, Sigma) and co-precipitating proteins were detected by western blot. Direct interactions were tested *in vitro* using recombinant proteins purified from *E. coli*. Protein interactions were detected by SDS-PAGE and Coomassie staining. Primer extensions were performed using RNA extracted from cells where RNPs were reconstituted without NP using a micro vRNA template (77 nt derived from *NP*) (Mondal et al., 2017; Turrell et al., 2013).

## Acknowledgements

Anti-influenza virus RNP antiserum (NR-3133) was obtained through NIH Biodefense and Emerging Infections Research Resources Repository, NIAID, NIH. We thank members of the Mehle lab for critical reading of the manuscript. This work was supported by NIH/NIAID R01AI125271 and the Greater Milwaukee Foundation Shaw Scientist Award to A.M. S.F.B. was supported by NIAID T32AI55397. A.M. is a Burroughs Wellcome Fund Investigator in the Pathogenesis of Infectious Disease.

## Author Contributions

Conceptualization, S.F.B. and A.M.; Methodology, S.F.B., M.P.L., and A.M.; Formal Analysis; S.F.B. and M.P.L.; Investigation, S.F.B.; Writing – Original Draft,

S.F.B. and A.M.; Writing – Review & Editing, S.F.B. and A.M.; Visualization, S.F.B. and A.M.; Funding Acquisition, S.F.B. and A.M.; Supervision, A.M.

